# FERONIA limits jasmonic acid overaccumulation and oxidative stress to enable plant survival at elevated temperatures

**DOI:** 10.64898/2026.05.21.727061

**Authors:** Jeonghyang Park, Jinwoo Park, Geonhee Hwang, Nayoung Lee, Eunkyoo Oh

**Affiliations:** Department of Life Sciences, Korea University, Seoul, Korea; Division of Applied Life Science (BK21four), Research Institute of Life Science, Gyeongsang National University, Jinju, Korea

## Abstract

Plants, as sessile organisms, must continually adapt to fluctuating temperatures to ensure survival. The plasma membrane-localized receptor-like kinase FERONIA (FER) coordinates diverse physiological processes and responses to various biotic and abiotic stresses. However, the role of FER in plant adaptation to elevated temperatures remains largely unexplored. Here, we report that FER is indispensable for plant thermotolerance. We found that *fer* loss-of-function mutants exhibit impaired thermomorphogenic growth and are hypersensitive to mild heat stress, displaying extensive oxidative stress-mediated cell death at elevated temperatures. Combined genetic and molecular analyses revealed that these temperature-sensitive defects in *fer* mutants are caused by an overaccumulation of jasmonic acid (JA), which subsequently triggers excessive production of reactive oxygen species. Furthermore, we show that this aberrant JA accumulation and oxidative stress are attributable to impaired FER-mediated regulation of turgor-dependent cell wall tensile stress. Taken together, our results suggest that FER-mediated cell wall tensile stress regulation serves as a critical mechanism to prevent aberrant JA accumulation and oxidative stress at elevated temperatures, thereby enabling plants to adapt to and survive under high-temperature conditions.

## Introduction

Plants, as sessile organisms, are constantly exposed to fluctuating environmental conditions and must adapt their physiology and development to ensure survival. Among the various abiotic stresses, elevated temperatures represent one of the most detrimental factors, severely impairing vegetative growth, reproductive success, and ultimately crop productivity. To mitigate the damaging effects of heat stress, plants have evolved multiple adaptive strategies, including morphological changes known as thermomorphogenesis, which enhance cooling capacity, and the rapid accumulation of heat-shock proteins (HSPs), which function as molecular chaperones to protect proteins from misfolding and aggregation (Casal and Balasubramanian, 2019; Dundar et al., 2025; Hwang et al., 2025). In addition, plants can acquire thermotolerance through heat acclimation, a process that reprograms gene expression, hormone signaling, and cellular metabolism to enhance survival under future heat stress conditions (Baurle, 2016).

FERONIA (FER) is a plasma membrane–localized receptor-like kinase belonging to the *Catharanthus roseus* Receptor-like Kinase 1-Like (CrRLK1L) family. Members of this family are characterized by an extracellular region containing two malectin-like domains, a single-pass transmembrane domain, and an intracellular kinase domain. FER was first identified for its essential role in pollen tube reception, a critical step in plant fertilization (Escobar-Restrepo et al., 2007). Subsequent studies have revealed that FER functions in a remarkably wide range of biological processes, including root hair development, morphogenesis of epidermal pavement cells, responses to abiotic stresses such as salt, diverse hormone responses, cell wall integrity maintenance, and immunity (Chaudhary et al., 2025; Deslauriers and Larsen, 2010; Duan et al., 2010; Feng et al., 2018; Lin et al., 2022; Liu et al., 2023a; Stegmann et al., 2017; Zhao et al., 2021; Zhao et al., 2018).

The remarkably diverse biological roles of FER are partly attributable to its ability to interact with components of many different signaling pathways, such as LRE-LIKE GPI-ANCHORED PROTEIN 1 (LLG1), LEUCINE RICH REPEAT EXTENSINS (LRXs), FLAGELLIN-SENSING 2 (FLS2), BRASSINOSTEROID INSENSITIVE 1-ASSOCIATED KINASE 1 (BAK1), ABA INSENSITIVE 2 (ABI2), BRASSINOSTEROID-INSENSITIVE 2 (BIN2), phytochrome B (phyB), and others (Chaudhary et al., 2025; Chen et al., 2016; Dunser et al., 2019; Liu et al., 2023b; Stegmann et al., 2017; Xiao et al., 2019). FER also binds RAPID ALKALINIZATION FACTOR (RALF) peptides in conjunction with its co-receptor LLG1 (Xiao et al., 2019). A recent study showed that RALF peptides, by interacting with the cell wall polysaccharide pectin, form pectin–RALF–FER–LLG1 condensates that trigger endocytosis of non-cognate regulators involved in diverse processes (Liu et al., 2024), providing a molecular basis for the pleiotropic functions of FER. In addition to RALFs, FER also directly binds to pectin through the malectin-like domains in its extracellular domain, and has been proposed to sense mechanical perturbations at the cell wall–plasma membrane interface arising from turgor-dependent changes in cell wall tensile stress (Feng et al., 2018; Lin et al., 2022; Malivert and Hamant, 2023; Qin et al., 2026). In *fer* loss-of-function mutants, the failure to sense and compensate for such mechanical perturbations frequently results in cell bursting under conditions of high cell wall tension, which has been proposed to underlie the pleiotropic defects observed in diverse biological processes (Malivert and Hamant, 2023).

In this study, we found that FER is required for plant thermotolerance. The *fer* mutants are hypersensitive to mild heat stress, resulting in oxidative stress-mediated cell death at elevated temperatures. We show that overaccumulation of jasmonic acid (JA) under high temperatures is responsible for the excessive accumulation of reactive oxygen species (ROS) and thereby for the reduced thermotolerance in *fer* mutants. We further show that the excessive accumulation of JA and ROS during heat stress is attributable to the disruption of FER-mediated cell wall tensile stress regulation. Together, our results establish that this regulatory mechanism is essential to prevent JA-induced oxidative stress, thereby enabling plants to tolerate heat stress.

## Results

### FER is required for tolerance to mild heat stress

*Arabidopsis thaliana* plants respond to elevated temperature through a suite of morphological changes collectively termed thermomorphogenesis. Since FER is involved in plant tolerance to several abiotic stresses, we examined whether it also contributes to thermomorphogenic responses. To this end, we grew both wild-type (WT) and *fer-4* mutant plants under two different ambient temperatures (20 °C and 28 °C). WT plants grown at 28 °C exhibited characteristic thermomorphogenic traits, including significantly longer hypocotyls (Figure 1A). In contrast, *fer-4* hypocotyls failed to elongate at 28 °C, instead, these mutants showed severe growth inhibition and leaf bleaching (Figure 1A). Trypan blue staining revealed that a significant portion of cotyledon and hypocotyl cells underwent cell death in *fer-4* mutants, but not in WT, when grown at 28 °C (Figure 1B). Propidium iodide (PI) staining further confirmed the occurrence of cell death, as evidenced by marked PI penetration into *fer-4* cells at 28 °C (Figure 1C). Taken together, these results indicate that the thermomorphogenic defects in *fer-4* are a consequence of cell death and that FER activity is critical for plant growth and survival at moderately elevated temperatures.

**Figure 1.**
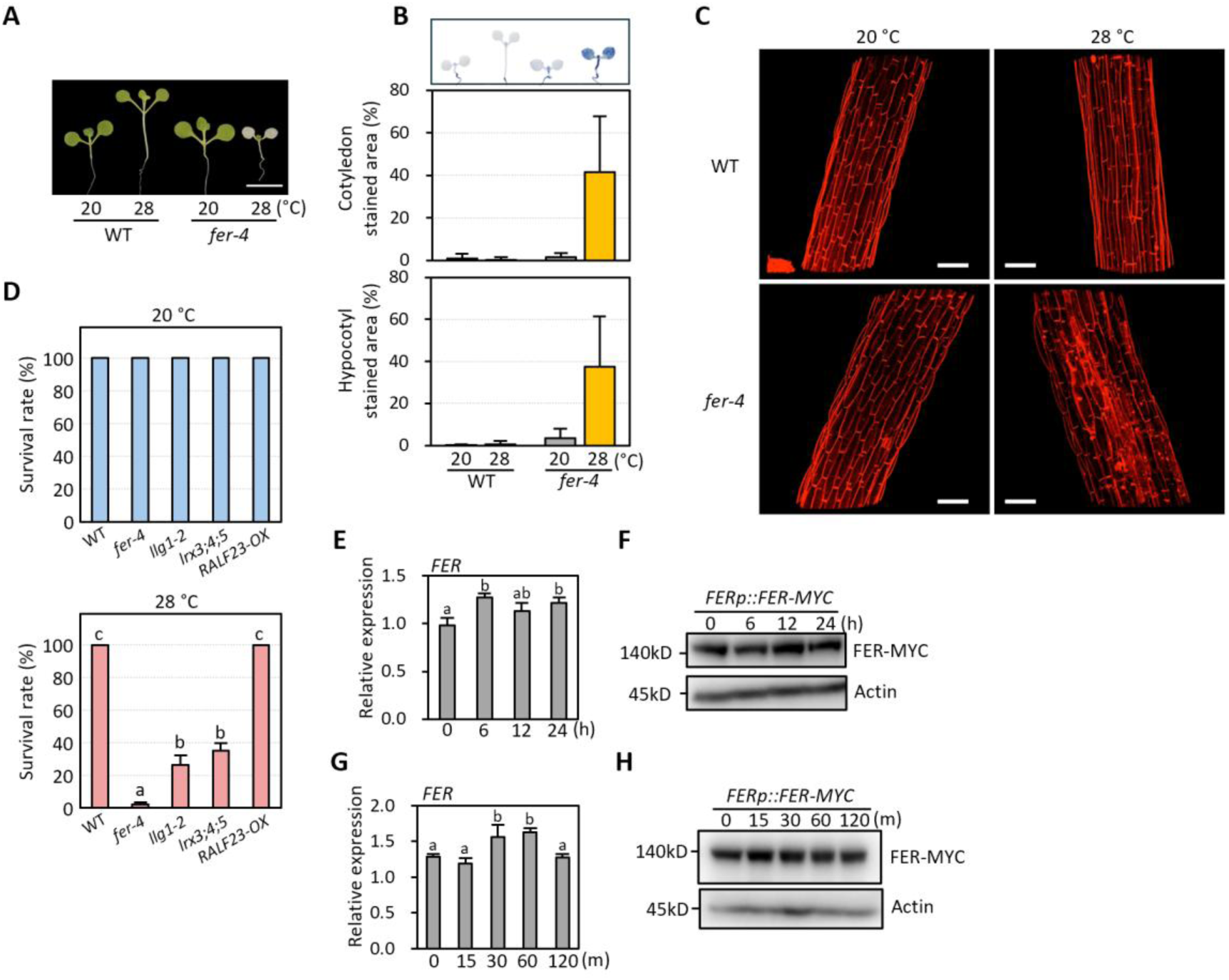
FER is required for tolerance to mild heat stress. **(A)** Representative images of wild-type (WT) and *fer-4* plants grown at two different temperatures. Plants were grown under continuous white light at 20 °C for 4 days and then transferred to either 20 or 28 °C for 6 days. **(B)** Trypan blue staining of WT and *fer-4* at 20 °C and 28 °C. Plants were grown at 20 °C for 4 days and then transferred to either 20 °C or 28 °C for 3 days before harvesting for staining. Representative images of plants are shown in the upper panel. Staining intensity was quantified using the ImageJ software. Letters above the bars indicate significant differences as determined by one-way ANOVA followed by Tukey’s test (*P* < 0.05). Error bars represent SD from at least fifteen biological replicates (*n* ≥ 15). **(C)** Propidium iodide (PI) staining of WT and *fer-4* mutants at 20 °C and 28 °C. Plants were grown at 20 °C for 4 days and then transferred to either 20 °C or 28 °C for 3 days before harvesting for staining. After staining, the plants were visualized using a confocal laser scanning microscope. Scale bars indicate 100 μm. **(D)** Survival rates of plants grown at 20 °C and 28 °C. Plants were initially grown at 20 °C for 4 days and then transferred to either 20 or 28 °C for 18 days before determining survival rates. Letters above the bars indicate significant differences determined by one-way ANOVA followed by Tukey’s test (*P* < 0.05). Error bars indicate SD from three independent biological replicates, each consisting of at least 70 seedlings. **(E and G)** qRT-PCR analysis of *FER*. Plants were grown at 20°C for 7 days and then transferred to 28 °C for the indicated time points before harvesting for total RNA extraction. Gene expression levels were normalized to *PP2A*. Letters above the bars indicate significant differences determined by one-way ANOVA followed by Tukey’s test (*P* < 0.05). Error bars represent SD from three independent biological replicates, each consisting of at least 40 seedlings. **(F and H)** Immunoblot analysis of FER-MYC proteins in plants grown under the same conditions as in (E) and (G). Immunoblotting was carried out using anti-MYC and anti-Actin (as a loading control) antibodies.

FER associates with its co-receptor LLG1 that functions as a chaperone for the proper membrane localization of FER and as a co-receptor for RALF peptide recognition (Li et al., 2015; Xiao et al., 2019). Both FER and RALFs interact with extracellular LRX proteins (Dunser et al., 2019; Li et al., 2015; Moussu et al., 2020; Zhao et al., 2021). Additionally, *llg1-2* and *lrx3;4;5* triple mutants exhibit morphological phenotypes similar to *fer-4* mutants. Therefore, we determined whether these FER-associated components are required for FER-mediated thermotolerance. Although the phenotype was less severe than that of *fer-4* mutants, both *llg1-2* and *lrx3;4;5* triple mutants exhibited severe leaf bleaching at 28 °C, but not at 20 °C (Figure 1D), suggesting that a signaling complex composed of FER, LLG1, and LRXs is essential for maintaining normal growth under moderately elevated temperatures (Figure 1D).

Given that LLG1 and the LRXs, along with RALF peptides, are known to mediate FER-dependent salt tolerance, we hypothesized that RALFs might also participate in FER-mediated thermotolerance. However, plants overexpressing RALF23 (*RALF23-OX*) showed no detectable defects in thermotolerance (Figure 1D), even though they were highly susceptible to salt stress (Zhao et al., 2018). This finding implies that FER regulates thermotolerance and salt tolerance through distinct signaling mechanisms.

We next examined whether *FER* expression or FER protein abundance is altered by temperature changes using *FER-MYC;fer-4* plants, in which MYC-fused FER is driven by its native promoter. Neither *FER* expression nor FER protein abundance was significantly affected by short-term exposure to high temperature (28 °C) (Figure 1E, F). Furthermore, prolonged exposure to high temperature did not significantly affect *FER* expression or protein levels (Figure 1G, H). These results indicate that FER protein abundance is not modulated by temperature changes.

### FER prevents excessive reactive oxygen species accumulation at high temperatures

To identify why *fer-4* mutants are susceptible to high temperature stress, we performed RNA sequencing (RNA-seq) analyses using WT and *fer-4* mutants grown continuously at 20 °C or exposed to 28 °C for 6 h. RNA-Seq analyses identified 1,477 and 2,271 high-temperature-regulated genes in WT and *fer-4* mutants, respectively (Figure 2A, B; Data S1). Among the 1,477 high-temperature-regulated genes in WT, 596 were activated, and 881 were repressed by exposure to 28 °C (Figure 2A, B). On the other hand, 1,114 genes were activated, and 1,157 genes were repressed by high temperature exposure in *fer-4* mutants (Figure 2A, B). The higher number of high-temperature-regulated genes in *fer-4* mutants is consistent with their high temperature susceptible phenotypes. Moreover, many genes that responded to high temperature in *fer-4* were not affected in WT (Figure 2A, B). Gene ontology (GO) enrichment analysis revealed that GO terms such as “jasmonic acid biosynthetic process”, “response to salt stress”, “response to salicylic acid”, “response to abscisic acid”, and “response to oxidative stress” were significantly enriched in *fer-4*-specific high-temperature-activated genes (Figure 2C). Conversely, GO terms related to DNA replication were enriched among the genes repressed by high temperature specifically in *fer-4* (Figure 2C).

**Figure 2.**
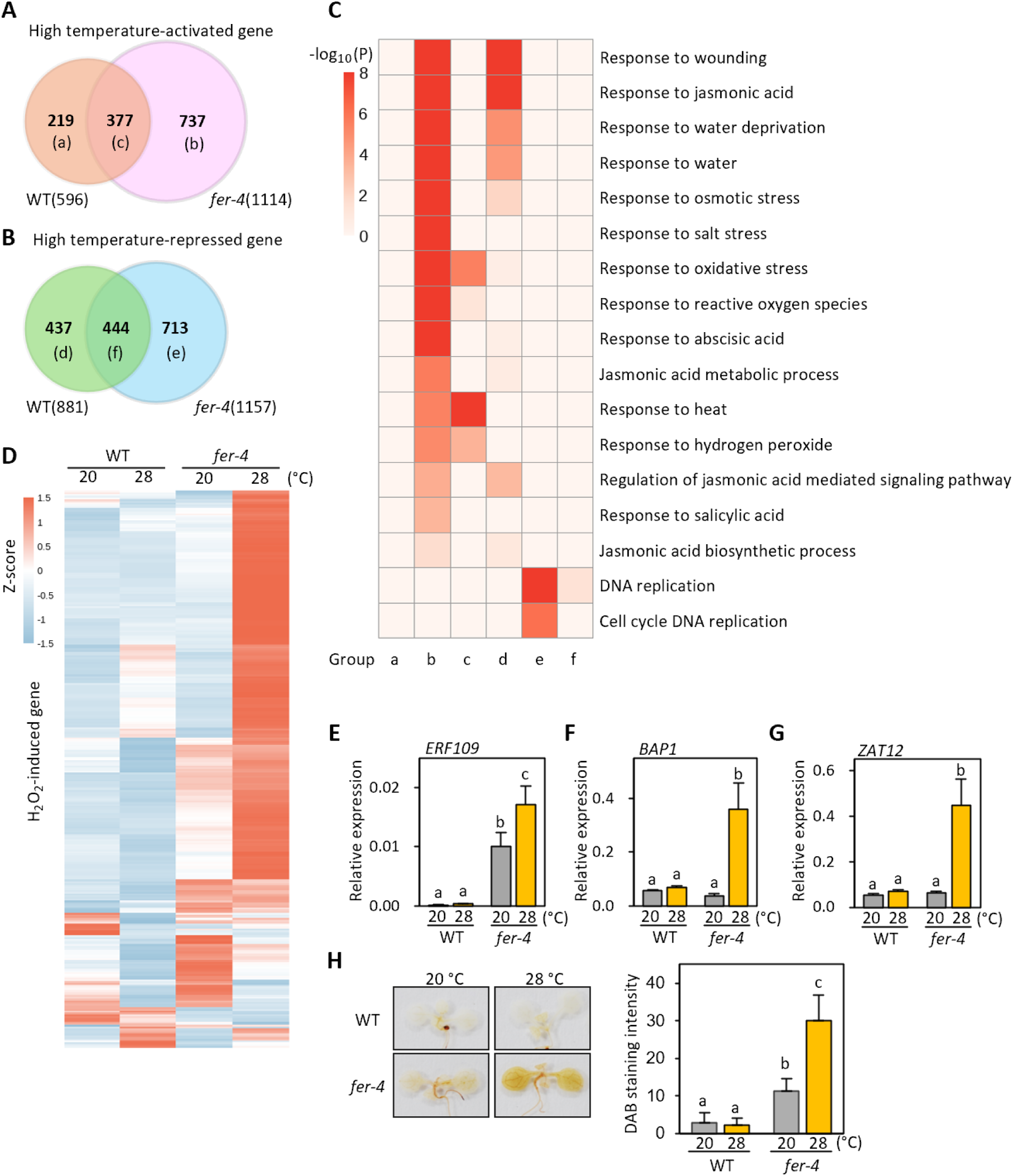
FER prevents excessive reactive oxygen species accumulation at high temperatures. **(A and B)** Venn diagrams showing the overlap of high-temperature-activated (A) and -repressed (B) genes between WT and *fer-4.* **(C)** Gene ontology (GO) enrichment analysis of high-temperature-regulated genes. The heatmap shows the enrichment of representative GO terms for the gene groups identified in (A) and (B). The color scale represents -log_10_(*P*). **(D)** Heatmap showing the expression profiles of H_2_O_2_-induced genes in WT and *fer-4* at 20 °C and 28 °C. Relative expression levels are presented as row-normalized z-scores derived from normalized TPM values from RNA-seq analysis. **(E–G)** qRT-PCR analysis of oxidative stress-induced genes *ERF109* (E), *BAP1* (F) and *ZAT12* (G). Plants were grown at 20 °C for 7 days and then transferred to either 20 °C or 28 °C for 24 h before harvesting for total RNA extraction. Gene expression levels were normalized to *PP2A*. Letters above the bars indicate significant differences determined by one-way ANOVA followed by Tukey’s test (*P* < 0.05). Error bars represent SD from three independent biological replicates, each consisting of at least 40 seedlings. **(H)** DAB staining of WT and *fer-4* plants at 20 °C and 28 °C. Staining intensity was quantified using the ImageJ software. Letters above the bars indicate significant differences as determined by one-way ANOVA followed by Tukey’s test (*P* < 0.05). Error bars represent SD from ten biological replicates (*n* = 10).

Among the genes activated by high temperature specifically in *fer-4*, we focused on those involved in oxidative stress responses, as these pathways are critical for abiotic stress tolerance. The expression of several oxidative stress-related genes, including those responsive to singlet oxygen and hydrogen peroxide (H₂O₂) (Davletova et al., 2005; op den Camp et al., 2003), was upregulated specifically in *fer-4* mutants under high temperature, whereas no such induction was observed in WT (Figure 2D and S1). Consistent with these results, quantitative RT-PCR (qRT-PCR) analyses confirmed similar expression patterns for three oxidative stress marker genes (*ERF109*, *BAP1*, and *ZAT12*) (Figure 2E-G).

We next performed 3,3′-diaminobenzidine (DAB) staining to determine reactive oxygen species (ROS) accumulation in *fer-4* mutants. DAB staining revealed stronger H₂O₂ accumulation in *fer-4* than in WT at 20 °C (Figure 2H). Notably, H₂O₂ levels remained largely unchanged in WT upon exposure to high temperature but increased markedly in *fer-4* (Figure 2H). These results suggest that excessive oxidative stress contributes, at least in part, to the reduced thermotolerance of *fer-4* mutants.

### Overaccumulation of jasmonic acid compromises thermotolerance in *fer-4* mutants

In addition to the “response to oxidative stress” GO term, terms such as “response to abscisic acid” and “response to salicylic acid” terms were also significantly enriched among the genes activated by high temperature specifically in *fer-4* mutants (Figure 2C). To determine whether enhanced abscisic acid (ABA) or salicylic acid (SA) responses contribute to the reduced thermotolerance of *fer-4*, we generated and analyzed *fer-4;aba2-1*, *fer-4;sid2*, and *fer-4;npr1* double mutants. However, genetic disruption of ABA biosynthesis, SA biosynthesis, or SA signaling failed to restore thermotolerance of *fer-4*, indicating that exaggerated SA or ABA responses are not responsible for its heat susceptibility (Figure 3A).

**Figure 3.**
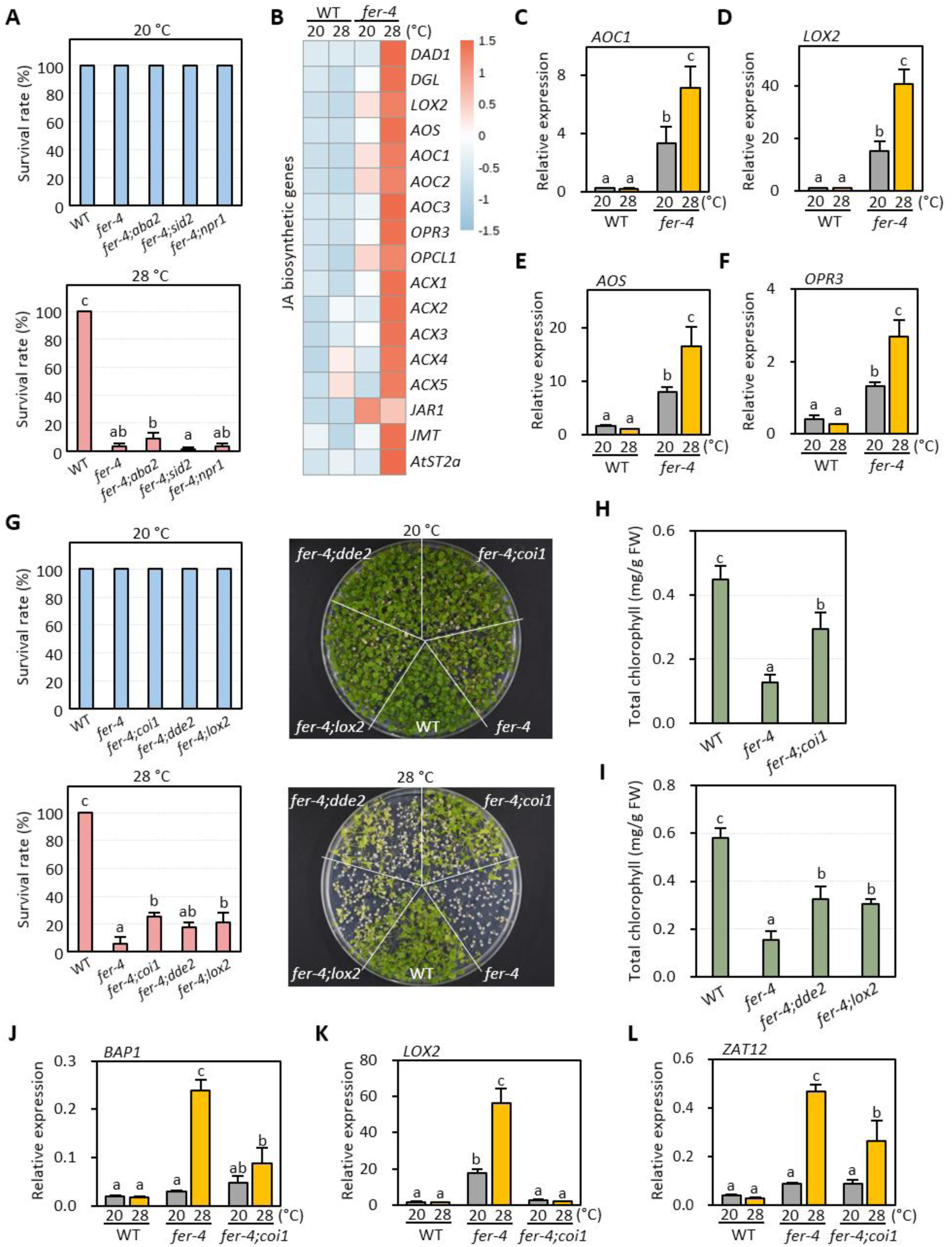
Overaccumulation of jasmonic acid compromises thermotolerance in *fer-4* mutants. **(A and G)** Survival rates of plants grown at 20 °C and 28 °C. Plants were initially grown at 20 °C for 4 days and then transferred to either 20 or 28 °C for 18 days before determining survival rates. Representative images of plants are shown in the right panels in (G). Letters above the bars indicate significant differences determined by one-way ANOVA followed by Tukey’s test (*P* < 0.05). Error bars indicate SD from three independent biological replicates, each consisting of at least 70 seedlings. **(B)** Heatmap showing the expression profiles of JA biosynthetic genes in WT and *fer-4* at 20 °C and 28 °C. Relative expression levels are presented as row-normalized z-scores derived from normalized TPM values from RNA-seq analysis. **(C–F)** qRT-PCR analysis of JA biosynthetic genes *AOC1* (C), *LOX2* (D), *AOS* (E) and *OPR3* (F). Plants were grown at 20 °C for 7 days and then transferred to either 20 °C or 28 °C for 24 h before harvesting for total RNA extraction. Gene expression levels were normalized to *PP2A*. Letters above the bars indicate significant differences determined by one-way ANOVA followed by Tukey’s test (*P* < 0.05). Error bars represent SD from three independent biological replicates, each consisting of at least 40 seedlings. **(H and I)** Chlorophyll accumulation in plants grown at 20 °C or 28 °C. Plants were grown under the same conditions as in (A) and (G). Error bars represent SD from seven independent biological replicates. Letters above the bars indicate significant differences determined by one-way ANOVA followed by Tukey’s test (*P* < 0.05). **(J–L)** qRT-PCR analysis of *BAP1* (J), *LOX2* (K) and *ZAT12* (L). Plants were grown at 20 °C for 7 days and then transferred to either 20 °C or 28 °C for 24 h before harvesting for total RNA extraction. Gene expression levels were normalized to *PP2A*. Letters above the bars indicate significant differences determined by one-way ANOVA followed by Tukey’s test (*P* < 0.05). Error bars represent SD from three independent biological replicates, each consisting of at least 40 seedlings.

The GO terms “Response to jasmonic acid” and “Jasmonic acid biosynthetic process” were also significantly enriched among the *fer-4*-specific high-temperature-activated genes (Figure 2C). At 20 °C, the expression of several jasmonic acid (JA) biosynthetic genes—including *DGL*, *LOX2*, *AOS*, *AOCs*, and *OPR3*—was higher in *fer-4* than in WT (Figure 3B). Intriguingly, upon exposure to high temperature, the expression of these genes further increased in *fer-4*, whereas no such induction was observed in WT (Figure 3B).

We confirmed these transcriptomic patterns by qRT-PCR analysis. In WT, the expression of *AOC1*, *LOX2*, *AOS*, and *OPR3* either decreased or remained unchanged following high-temperature treatment (Figure 3C-F). In contrast, all these genes were transcriptionally upregulated in *fer-4* under the same conditions (Figure 3C-F). These results indicate that high temperature enhances JA biosynthesis specifically in *fer-4*, but not in WT.

It has been reported that JA-deficient mutants display increased tolerance to high light stress (Ramel et al., 2013b). Based on this, we hypothesized that excessive JA accumulation contributes to the reduced thermotolerance of *fer-4*. To test this, we crossed *fer-4* with JA-signaling and JA-deficient mutants to generate *fer-4;coi1-16*, *fer-4;dde2*, and *fer-4;lox2* double mutants. Supporting our hypothesis, the survival rate of *fer-4;coi1-16* at 28 °C was significantly higher than that of *fer-4* (Figure 3G). Additionally, *fer-4;coi1-16* exhibited 2.3-fold higher chlorophyll content compared to *fer-4* at 28 °C (Figure 3H). Similarly, both *fer-4;dde2* and *fer-4;lox2* plants displayed improved thermotolerance (Figure 3G, I). Consistent with these phenotypes, the high-temperature-induced expression of JA biosynthetic and oxidative stress genes was attenuated in *fer-4;coi1-16* compared to *fer-4* (Figure 3J-L). Together, these results demonstrate that overaccumulation of JA under elevated temperature conditions contributes, at least in part, to the heat-susceptible phenotype of *fer-4* mutants.

### Both direct pectin binding and kinase activity are required for FER-mediated thermotolerance

The extracellular domain of FER contains tandem malectin-like domains A and B (MALA and MALB). Previous studies have shown that the MALA domain mediates FER binding to pectin, enabling it to monitor cell wall tensile stress at the cell wall–plasma membrane interface (Feng et al., 2018; Lin et al., 2022; Qin et al., 2026). To determine whether this pectin binding is required for FER-dependent thermotolerance, we tested the ability of FER variants to complement the *fer-4* mutant. Expression of FER-MYC fully restored thermotolerance in *fer-4*, whereas FER^ΔMALA^-MYC failed to do so (Figure 4A). Consistent with this result, the expression of JA biosynthetic and oxidative stress genes was restored to WT levels by the introduction of FER-MYC, but not FER^ΔMALA^-MYC, at 28 °C (Figure 4B-D). The protein levels of FER variants in *FER^ΔMALA^-MYC;fer-4* plants were comparable to those of FER in *FER-MYC;fer-4* plants (Figure 4E). Therefore, these results indicate that pectin binding through the MALA domain is essential for FER-mediated thermotolerance.

**Figure 4.**
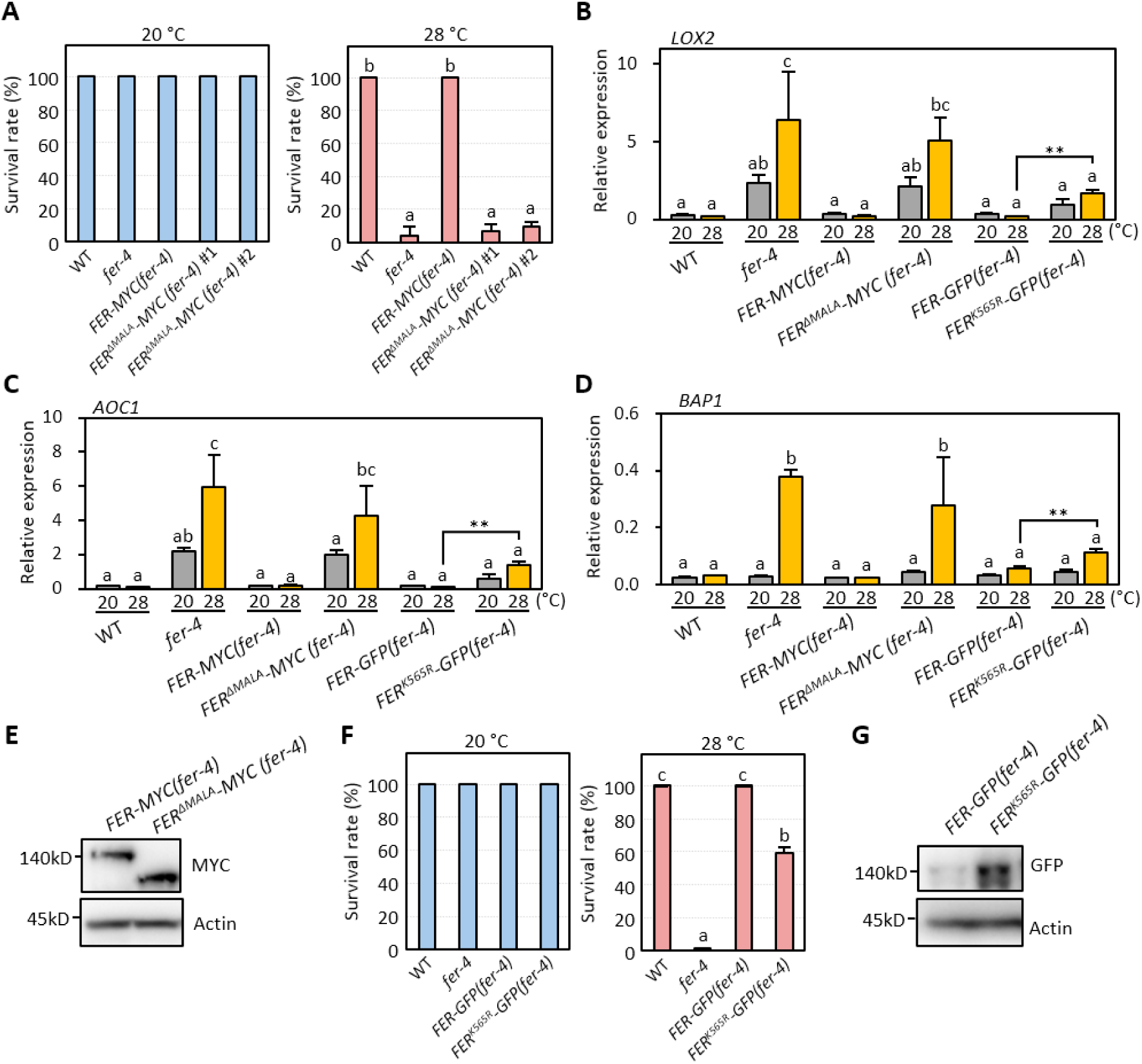
Both direct pectin binding and kinase activity are required for FER-mediated thermotolerance. **(A and F)** Survival rates of plants grown at 20 °C and 28 °C. Plants were initially grown at 20 °C for 2 days and then transferred to either 20 or 28 °C for 6 days before determining survival rates. Scale bar indicates 5 mm. Letters above the bars indicate significant differences determined by one-way ANOVA followed by Tukey’s test (*P* < 0.05). Error bars indicate SD from three independent biological replicates, each consisting of at least 70 seedlings. **(B–D)** qRT-PCR analysis of *LOX2* (B), *AOC1* (C) and *BAP1* (D). Plants were grown at 20 °C for 7 days and then transferred to either 20 °C or 28 °C for 24 h before harvesting for total RNA extraction. Gene expression levels were normalized to *PP2A*. Letters above the bars indicate significant differences determined by one-way ANOVA followed by Tukey’s test (*P* < 0.05). Error bars represent SD from three independent biological replicates, each consisting of at least 40 seedlings. **(E)** Immunoblot analysis of FER-MYC and FER^ΔMALA^-MYC proteins in plants grown at 20 °C for 7 days. Immunoblotting was carried out using anti-MYC and anti-Actin (as loading control) antibodies. **(G)** Immunoblot analysis of FER-GFP and FER^K565R^-GFP proteins in plants grown at 20 °C for 7 days. Immunoblotting was carried out using anti-GFP and anti-Actin (as loading control) antibodies.

We next investigated whether FER kinase activity is required for thermotolerance. While FER-GFP fully restored thermotolerance in *fer-4* mutants, a kinase-dead variant, FER^K565R^-GFP, only partially restored thermotolerance (Figure 4F). Consistently, the expression of *LOX2*, *AOC1*, *BAP1* was less effectively suppressed by FER^K565R^-GFP than by FER-GFP (Figure 4B-D). Given that FER levels were much higher in *FER^K565R^-GFP;fer-4* plants than in *FER-GFP;fer-4* plants (Figure 4G), these results suggest that FER kinase activity is critical for full thermotolerance, although FER retains some function independent of its kinase activity.

### Impaired FER-mediated cell wall tensile stress regulation underlies the reduced thermotolerance of *fer-4* mutants

FER has been proposed to monitor turgor-dependent cell wall tensile stress at the cell wall–plasma membrane interface, and *fer* mutants exhibit cell bursting under conditions of high wall tension (Malivert et al., 2021; Malivert and Hamant, 2023; Qin et al., 2026). The pleiotropic phenotypes of *fer* mutants have been suggested to stem from this failure to sense and compensate for changes in turgor-dependent cell wall tensile stress (Malivert and Hamant, 2023). Furthermore, considering that the MALA domain, which mediates FER association with cell wall pectin and is required for cell wall-anchored turgor sensing (Feng et al., 2018; Lin et al., 2022; Qin et al., 2026), is essential for FER-mediated thermotolerance, it is likely that the high-temperature susceptibility of *fer-4* arises from increased cell wall tensile stress resulting from impaired FER-mediated stress regulation.

To test this hypothesis, we compared transcriptomic changes observed in *fer-4* with those induced by treatment with isoxaben (ISX), a cellulose synthase inhibitor that reduces cell wall mechanical bearing capacity by blocking new cellulose deposition during cell growth (Zhai et al., 2024). Of the 522 genes upregulated in *fer-4* compared to WT at 20 °C, 362 (∼69.3%) overlapped with ISX-activated genes (Figure 5A), confirming constitutive cell wall tensile stress in *fer-4*. Notably, a significant number of genes (438) specifically upregulated by high temperature in *fer-4* also overlapped with ISX-induced genes (Figure 5B), whereas few WT-specific high-temperature-activated genes (11) showed such overlap (Figure 5C). These results suggest that cell wall tensile stress is further exacerbated at high temperatures in *fer-4*, potentially accounting for its reduced thermotolerance. In line with this notion, both JA biosynthetic and oxidative stress genes were upregulated 4.0- to 8.8-fold by high temperature in ISX-treated plants, similar to *fer-4* mutants (Figure 5D-G). Consistent with these gene expression patterns, ISX-treated plants were more susceptible to high temperatures compared to non-treated plants (Figure 5H).

**Figure 5.**
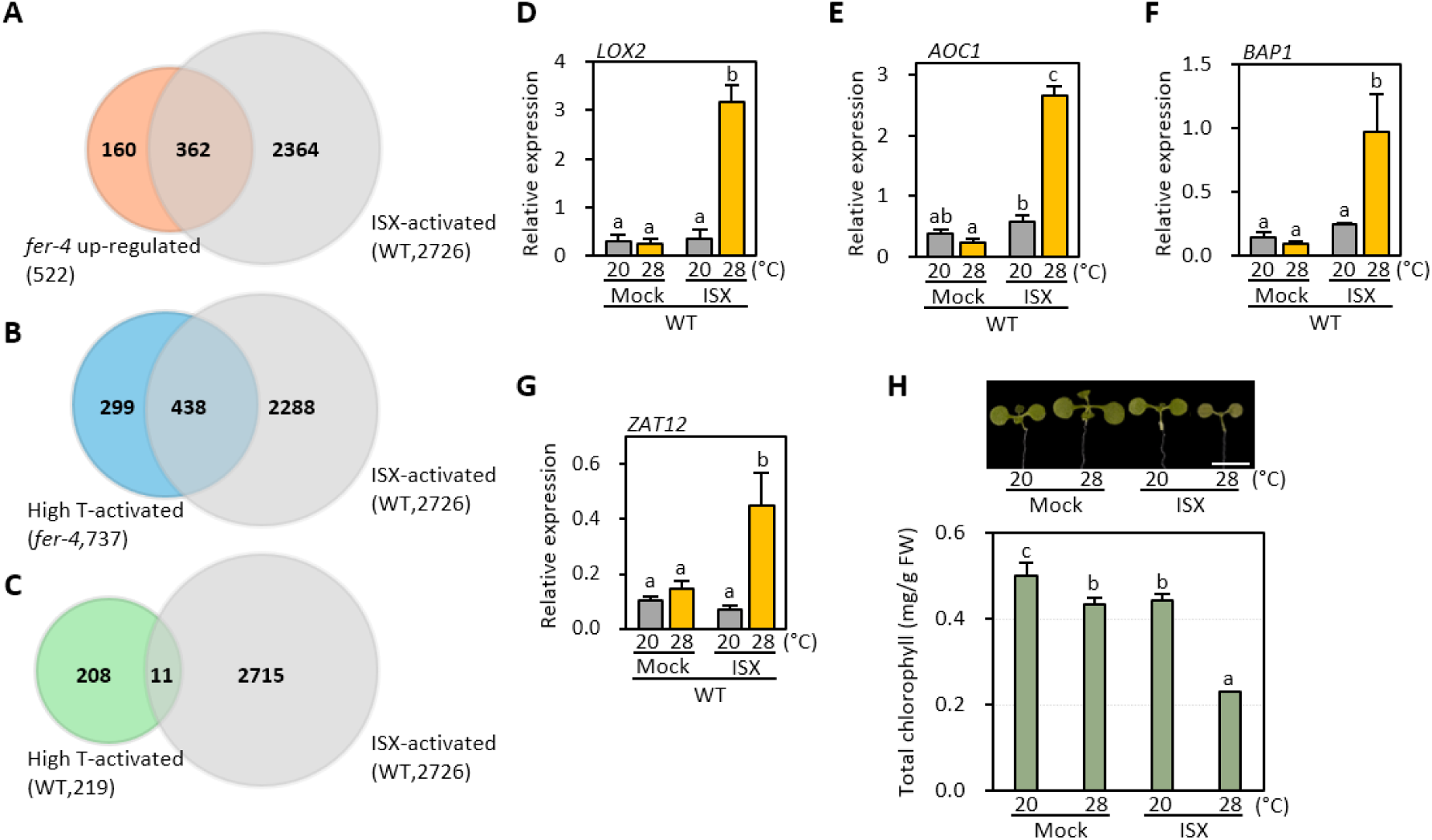
Cell wall perturbation by ISX compromises thermotolerance. **(A–C)** Venn diagrams showing the overlap of ISX-activated genes with genes up-regulated genes in *fer-4* relative to WT (A), high-temperature-activated genes specifically in *fer-4* (B), and high-temperature-activated genes in WT (C). The overlaps in (A) (362 genes) and (B) (438 genes) were highly significant (*P* < 10^−10^; *OR* = 28.57 and 19.03, respectively), whereas the overlap in (C) (11 genes) was not statistically significant (*P* > 0.05; *OR* = 0.58). Statistics were calculated using hypergeometric tests for enrichment. **(D–G)** qRT-PCR analysis of *LOX2* (D), *AOC1* (E), *BAP1* (F) and *ZAT12* (G). Plants were grown at 20 °C for 7 days and then transferred to liquid media supplemented with or without 10 nM ISX. The plants were subsequently incubated at either 20 °C or 28 °C for 24 h before harvesting for total RNA extraction. Gene expression levels were normalized to *PP2A*. Letters above the bars indicate significant differences determined by one-way ANOVA followed by Tukey’s test (*P* < 0.05). Error bars represent SD from three independent biological replicates, each consisting of at least 40 seedlings. **(H)** Chlorophyll accumulation in plants grown at 28 °C. WT and *fer-4* were grown at 20 °C for 7 days and then transferred to liquid media supplemented with mock and 10 nM ISX. The plants were subsequently incubated at 28 °C for 3 days before harvesting for the determination of chlorophyll content. The representative images of plants are shown in an upper panel. Scale bar indicates 5 mm. Letters above the bars indicate significant differences determined by one-way ANOVA followed by Tukey’s test (*P* < 0.05). Error bars represent SD from three independent biological replicates.

To further confirm the importance of FER-mediated cell wall tensile stress regulation in thermotolerance, we examined whether the survival rate of *fer-4* mutants at 28 °C could be improved by reducing cell wall tensile stress through altering the agar concentration of the growth medium. Increasing the agar concentration raises the osmolarity of the medium, thereby reducing turgor-dependent cell wall tensile stress. On the medium containing 0.7% agar, *fer-4* mutants were severely susceptible to high-temperature stress. In contrast, increasing the agar concentration to 2.5% largely restored thermotolerance (Figure 6A). Consistent with this observation, the induction of JA biosynthetic and oxidative stress genes was markedly attenuated in *fer-4* grown on 2.5% agar medium compared to those grown on 0.7% agar medium (Figure 6B-G). Notably, high temperature-induced hypocotyl elongation and the expression of auxin-responsive genes (*IAA19* and *YUC8*) were also restored in *fer-4* mutants when grown on 2.5% agar medium (Figure 6H and S2).

**Figure 6.**
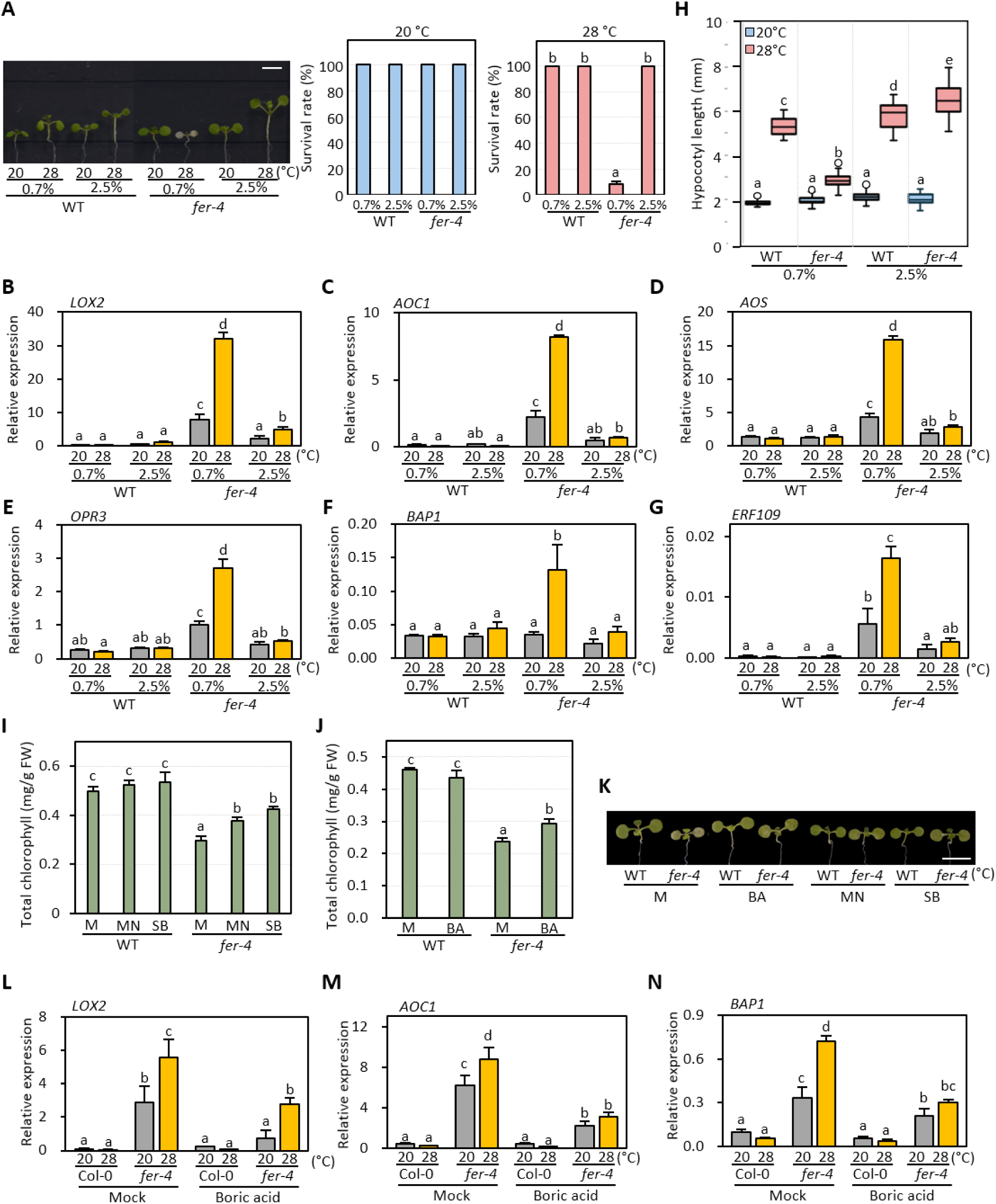
Increased cell wall tension underlies the reduced thermotolerance of *fer-4* mutants. **(A)** Survival rates of plants grown at 20 °C and 28 °C on different agar concentrations. WT and *fer-4* plants were grown at 20 °C for 2 days on the medium containing either 0.7% or 2.5% agar and then transferred to either 20 °C or 28 °C for 6 days before determining survival rates. The representative images of plants are shown in the left panel. Scale bar indicates 5 mm. Letters above the bars indicate significant differences determined by one-way ANOVA followed by Tukey’s test (*P* < 0.05). Error bars indicate SD from three independent biological replicates, each consisting of at least 70 seedlings. **(B–G)** qRT-PCR analysis of *LOX2* (B), *AOC1* (C), *AOS* (D), *OPR3* (E), *BAP1* (F) and *ERF109* (G). WT and *fer-4* plants were grown at 20 °C for 4 days on the medium containing either 0.7% or 2.5% agar and then either maintained at 20 °C or transferred to 28 °C for 24 h before harvesting for total RNA extraction. Gene expression levels were normalized to *PP2A*. Letters above the bars indicate significant differences determined by one-way ANOVA followed by Tukey’s test (*P* < 0.05). Error bars represent SD from three independent biological replicates, each consisting of at least 40 seedlings. **(H)** Effect of medium agar concentration on the hypocotyl elongation of plants. WT and *fer-4* plants were grown at 20 °C for 4 days on the medium containing either 0.7% or 2.5% agar and then transferred to either 20 °C or 28 °C for 3 days. Letters above the boxes indicate significant differences as determined by one-way ANOVA followed by Tukey’s test (*P* < 0.05). **(I–K)** Chlorophyll accumulation in plants grown at 28 °C. WT and *fer-4* were grown at 20 °C for 7 days and then transferred to liquid media supplemented with mock, 140 mM mannitol (MN) (I), 140 mM sorbitol (SB) (I), or 10 mM boric acid (BA) (J). The plants were subsequently incubated at 28 °C for 3 days before harvesting for the determination of chlorophyll content. The representative images of plants are shown in (K). Scale bar indicates 5 mm. Letters above the bars indicate significant differences determined by one-way ANOVA followed by Tukey’s test (*P* < 0.05). Error bars represent SD from three independent biological replicates. **(L–N)** qRT-PCR analysis of *LOX2* (L), *AOC1* (M), and *BAP1* (N). WT and *fer-4* were grown at 20 °C for 7 days and then transferred to liquid media supplemented with or without 10 mM boric acid. The plants were subsequently incubated at either 20 °C or 28 °C for 24 h before harvesting for total RNA extraction. Gene expression levels were normalized to *PP2A*. Letters above the bars indicate significant differences determined by one-way ANOVA followed by Tukey’s test (*P* < 0.05). Error bars represent SD from three independent biological replicates, each consisting of at least 40 seedlings.

Similarly, reducing cell wall tension by adding mannitol or sorbitol to the growth medium alleviated leaf bleaching in *fer-4* at 28 °C (Figure 6I, J). Furthermore, supplementing the medium with borate, which reduces cell wall tensile stress by reinforcing the cell wall through facilitating pectin cross-linking (Feng et al., 2018), mitigated the induction of *LOX2*, *AOC1*, and *BAP1* in *fer-4* mutants grown at 28 °C, and partially restored thermotolerance (Figure 6J-N). Together, these findings suggest that FER-mediated regulation of cell wall tensile stress is critical for thermotolerance.

Previous studies have established that FER is essential for maintaining cell wall integrity during brassinosteroid (BR)-mediated cell expansion (Chaudhary et al., 2025). Given that elevated temperatures activate BR signaling to promote cell elongation (Ibanez et al., 2018; Oh et al., 2012), we investigated whether this high temperature-activated BR signaling is responsible for the cell death observed in *fer-4* mutants. To test this, we treated seedlings with propiconazole (PPZ), a BR biosynthesis inhibitor, to reduce endogenous BR levels. However, the thermotolerance of *fer-4* was not restored even when BR biosynthesis was blocked and growth was significantly inhibited (Fig. S3), suggesting that the impaired thermotolerance of *fer-4* is independent of BR-mediated cell elongation.

## Discussion

Plants generally grow and develop within a specific temperature range, known as ambient temperature, without displaying stress-associated phenotypes (Wigge, 2013). WT *Arabidopsis thaliana* plants grow normally between 16 °C and 28 °C. However, in this study, we found that *fer-4* mutant plants exhibited severely retarded growth and leaf bleaching at 28°C, a temperature that does not affect WT plants. Furthermore, oxidative stress-responsive genes were highly induced, and ROS accumulated to high levels in *fer-4* mutants grown at 28°C, whereas WT plants remained unaffected. These results suggest that FER is required to prevent excessive ROS accumulation and ensure normal growth under mildly elevated temperatures.

Previous studies have established that FER contributes to tolerance against various abiotic stresses, particularly salt stress. Mutants defective in FER-associated components, such as *llg1-2* and *lrx3;4;5*, are highly susceptible to salt stress (Feng et al., 2018; Zhao et al., 2021; Zhao et al., 2018), indicating that FER regulates salt tolerance via these interacting partners. Interestingly, we found that *llg1-2* and *lrx3;4;5* mutants were also susceptible to high-temperature stress, exhibiting phenotypes similar to *fer-4*. This suggests that FER mediates thermotolerance through these same components. However, while the overexpression of RALF23 markedly reduced salt tolerance (Zhao et al., 2021; Zhao et al., 2018), it did not significantly affect thermotolerance. Furthermore, disruption of ABA biosynthesis restored salt tolerance in *fer-4* and *lrx3;4;5* mutants but failed to rescue thermotolerance in *fer-4* (Zhao et al., 2021). Collectively, these findings imply that while FER regulates both salt and high-temperature responses through common interacting components, such as LLG1 and LRXs, the underlying downstream signaling mechanisms are distinct.

Our RNA-seq analysis revealed that JA biosynthetic genes were strongly induced by elevated temperatures in *fer-4* mutants but not in WT. Notably, disruption of JA perception in *fer-4* mutants compromised the induction of oxidative stress genes under high temperatures. Consistently, mutations in either JA biosynthesis or perception partially restored thermotolerance and suppressed high-temperature-induced leaf bleaching in *fer-4* mutants. Previous studies have shown that excessive JA accumulation induces ROS production, subsequently leading to cell death (Ramel et al., 2013a; Rao et al., 2000; Zhang and Xing, 2008). Thus, our results suggest that FER enhances thermotolerance by restricting JA accumulation at high temperatures, thereby preventing excessive ROS production.

Bioactive JA accumulation is reduced at high temperatures due to enhanced JA catabolic pathways, a process required for thermomorphogenesis (Zhu et al., 2021). However, JA biosynthetic gene expression was increased by high temperature in *fer-4* mutants. Since JA biosynthesis is known to be upregulated in response to cell wall tensile stress (Ellis et al., 2002; Mielke et al., 2021), it is plausible that in the absence of FER, cell wall tensile stress is further exacerbated under high-temperature conditions.

We propose that in *fer-4* mutants, FER-mediated regulation of cell wall tensile stress is impaired, rendering cells unable to maintain cell wall mechanical homeostasis under high-temperature conditions. Our study provides several lines of evidence supporting this hypothesis. First, the MALA domain, which mediates FER binding to pectin and is required for cell wall tensile stress sensing (Feng et al., 2018; Lin et al., 2022; Qin et al., 2026), was essential for FER-mediated thermotolerance. Second, WT plants treated with ISX, a cellulose biosynthesis inhibitor that progressively reduces cell wall mechanical bearing capacity during cell growth, showed increased activation of JA biosynthesis and oxidative stress genes at high temperatures and became susceptible to mild heat stress. Third, borate treatment, which reinforces the cell wall through pectin cross-linking and thereby reduces tensile stress, mitigated JA biosynthetic and oxidative stress gene induction and partially restored thermotolerance in *fer-4* mutants. Finally, growing plants on high-agar medium, which reduces turgor-dependent cell wall tensile stress (Malivert and Hamant, 2023; Verger et al., 2018), suppressed JA biosynthetic and oxidative stress gene expression and restored thermotolerance in *fer-4* mutants.

Collectively, these results suggest that high temperature exacerbates cell wall tensile stress in *fer-4* mutants beyond a critical threshold, triggering JA overaccumulation and oxidative stress-mediated cell death. We therefore propose that FER-mediated regulation of cell wall tensile stress serves as a critical surveillance mechanism to prevent excessive JA and ROS accumulation at high temperatures, thereby enabling plants to grow and develop under mild heat stress.

While our research was ongoing, a recent study reported that FER regulates thermomorphogenesis by phosphorylating and inhibiting the non-canonical AUX/IAA protein IAA29 (Zheng et al., 2026). However, our data suggest a different primary cause for the thermomorphogenic growth defects in *fer-4* mutants. Although we also observed the lack of hypocotyl growth promotion in *fer-4* mutants at 28°C, the simultaneous occurrence of severe cell death in both hypocotyls and cotyledons and leaf bleaching suggests that the failure of thermomorphogenesis is a secondary consequence of loss of cell viability rather than a direct impairment of auxin signaling. In support of this notion, both thermomorphogenic responses and the expression of auxin-responsive genes in *fer-4* mutants were restored simply by increasing the agar concentration in the medium, a condition that prevents cell bursting by reducing turgor-dependent cell wall tensile stress.

Cell wall strength is largely determined by the degree of pectin methylesterification. It has been reported that pectin methylesterase (PME) activity is enhanced at high temperatures in soybean, leading to demethylesterified pectin that interacts more strongly with calcium ions, thereby strengthening the cell wall (Huang et al., 2017; Wu et al., 2010). Notably, a recent study demonstrated that PME activity is regulated by FER (Biermann et al., 2025). Therefore, it would be interesting to investigate in future studies whether FER regulates the level of pectin esterification to modulate cell wall tensile stress under high-temperature conditions.

## Materials and Methods

### Plant materials and growth conditions

For general growth and seed harvesting, *Arabidopsis thaliana* plants were grown in a growth room under 16/8 h light/dark cycles at 22 °C. All *A. thaliana* plants used in this study were the Col-0 ecotype background. The lines *fer-4*, *llg1-2*, *lrx3;4;5*, *RALF23-OX*, *FERp::FER-MYC;fer-4*, *FERp::FER-GFP;fer-4*, *FERp::FER^K565R^-GFP;fer-*4, *cop1-16*, and *dde2*-2 have been previously described (Chakravorty et al., 2018; Draeger et al., 2015; Shin et al., 2021; Srivastava et al., 2009; Xiao et al., 2019). The construct for generating *FERp::FER-MYC^ΔMALA^;fer-4* was generated using the primers listed in Table S1.

### Thermotolerance assay

To determine the survival rate, the degree of leaf bleaching was used as the criterion. Seedlings with more than 90% bleaching were classified as dead, while the remaining individuals were counted as survivors.

### Protein extraction and western blot analysis

Total protein was extracted using an extraction buffer (125mM Tris-HCl pH 6.8, 20% glycerol, 4% sodium dodecyl sulfate, 0.03mM bromophenol blue and 2% β-mercaptoethanol). FER-GFP and FER-MYC protein levels were detected by western blot analysis using anti-GFP (JL-8, 1:2000; Takara Bio) and anti-MYC (9B11, 1:2000; Cell Signaling Technology) antibodies, respectively. Actin protein levels were detected using an anti-actin antibody (AS13 2640, 1:5000; Agrisera) as a loading control.

### Quantitative real-time polymerase chain reaction (qRT-PCR) gene expression analysis

Total RNA was extracted from seedlings grown under specified conditions using the EasyPure Universal Plant Total RNA Kit (TransGen Biotech) according to the manufacturer’s instructions. First-strand cDNA synthesis was performed using M-MLV Reverse Transcriptase (Thermo Fisher Scientific). qRT-PCR was conducted on a CFX96 Real-Time PCR detection system (Bio-Rad) using an EvaGreen master mix (Solgent). Transcript levels of each gene were normalized to the internal reference gene *PP2A*. The experiments were carried out with three biological replicates, each consisting of pooled samples of over 40 seedlings. Gene-specific primers are listed in Table S1.

### Chlorophyll measurement

Approximately 20 mg of plant tissue grown under the specified conditions was harvested for total chlorophyll measurement. The samples were extracted with 0.5 mL of 95% (v/v) ethanol and incubated at 4°C overnight in the dark. After incubation, the absorbance at 645 nm and 663 nm was measured using a multi-mode reader (BioTek Synergy HTX, BioTek). Total chlorophyll content (mg/g fresh weight) was calculated using the following formula:

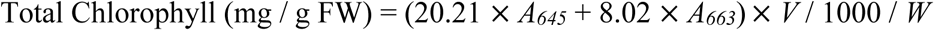

where *A_645_* and *A_663_* represent the absorbance at 645 nm and 663 nm, respectively; V is the total volume of the extract (mL); and *W* is the fresh weight of the sample (g).

### DAB staining

Hydrogen peroxide (H_2_O_2_) was visualized by DAB staining as described by (Daudi and O’Brien, 2012), with minor modification. Briefly, seedlings were vacuum-infiltrated with 1 mg/mL DAB staining solution and incubated for 3 h. Following staining, the seedlings were decolorized in 70% (v/v) ethanol at room temperature to remove chlorophyll. The ethanol was replaced once after 2 h, and the seedlings were kept overnight.

### PI staining and confocal microscopy

To visualize cell death, seedlings were subjected to propidium iodide (PI) staining. Seedlings were incubated in a 0.2 mg/mL PI solution (Sigma-Aldrich) for 7 min under dark conditions. Following incubation, the seedlings were washed two times with triple-distilled water. The stained samples were immediately mounted and visualized using a LSM800 confocal laser scanning microscope (ZEISS) equipped with an appropriate filter set for PI fluorescence (excitation at 535 nm and emission at 617 nm).

### RNA-seq analysis

Plants were grown under continuous white light at 20 °C for 7 days, followed by treatment at 20 °C or 28 °C for 6 h, and then harvested for RNA extraction. Total RNA was extracted using the MiniBEST Plant RNA Extraction Kit (Takara) according to the manufacturer’s instructions. mRNA sequencing libraries were constructed using the TruSeq RNA Sample Preparation Kit (Illumina) and sequenced on an Illumina NextSeq 500 platform. Transcript abundances were quantified against the TAIR10 transcriptome using Salmon (Patro et al., 2017), and differential expression analysis was performed using the DESeq2 R package. Differentially expressed genes (DEGs) were defined based on the following criteria: fold change ≥ 2 and adjusted *P*-value < 0.01 (Love et al., 2014). For visualization, normalized read counts of selected genes generated by DESeq2 using the median-of-ratios method were row-scaled (Z-score) and visualized as heatmaps using the *pheatmap* package in R. The RNA-seq data generated in this study have been deposited in the NCBI Gene Expression Omnibus (GEO) under accession number GSE317106.

## Author Contributions

Je.P., Ji.P., G.H., and N.L. designed and performed the experiments and analyzed the data. E.O. supervised the project and conceived the study, designed the experiments, analyzed the data, and wrote the manuscript with inputs from all the authors.

## Acknowledgments

We thank Sarah M Assmann for providing *FERp::FER-GFP;fer-4* and *FERp::FER^K565R^-GFP;fer-4* seeds, Hyo-Jun Lee for providing *FERp::FER-MYC;fer-4* seeds, Christoph Ringli for providing *lrx3;4;5* seeds, and Stephen H Howell for providing *RALF23-OX* seeds.

