## Supplementary material for "FERONIA limits jasmonic acid overaccumulation and oxidative stress to enable plant survival at elevated temperatures": Figure S1, S2, S3

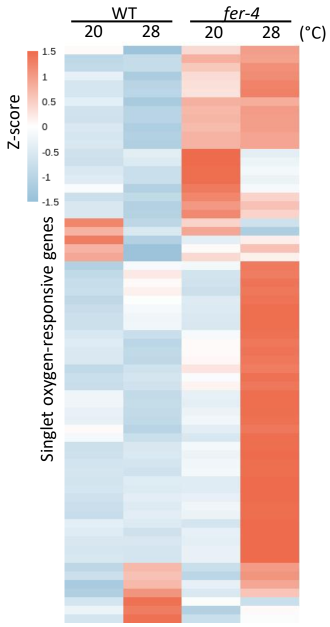


**Figure S1. High temperature activates singlet oxygen-responsive genes in *fer-4* mutants**

Heatmap showing the expression profiles of singlet oxygen-responsive genes in WT and *fer-4* at 20 °C and 28 °C. Relative expression levels are presented as row-normalized z-scores derived from normalized TPM values from RNA-seq analysis.

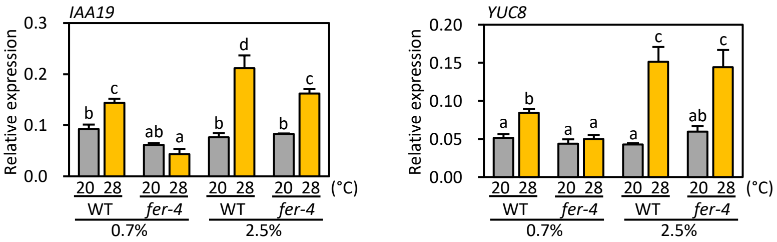


**Figure S2. Effect of medium agar concentration on the high temperature-activated auxin responsive gene expression in WT and *fer-4* mutants.**

qRT-PCR analysis of *IAA19* and *YUC8* expression. Plants were grown at 20°C for 7 days on the medium containing either 0.7% or 2.5% agar and then transferred to 20°C or 28 °C for 24 hours before harvesting for total RNA extraction. Gene expression levels were normalized to *PP2A*. Different letters above the bars indicate significant differences determined by one-way ANOVA followed by Tukey's test ($P$ < 0.05). Error bars represent SD from three independent biological replicates, each consisting of at least 40 seedlings.

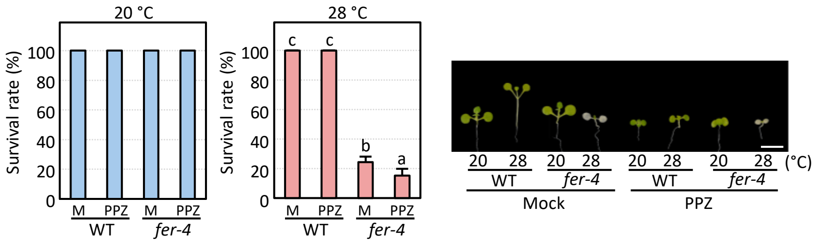


**Figure S3. The reduced thermotolerarnce of *fer-4* is independent of BR-mediated cell elongation.**

WT and *fer-4* plants were grown at 20 °C for 4 days on the medium supplemented with or without 0.2 $\mu$M PPZ and then transferred to either 20 °C or 28 °C for 6 days before determining survival rates. The representative images of plants are shown in the right panel. Scale bar indicates 5 mm. Letters above the bars indicate significant differences determined by one-way ANOVA followed by Tukey's test ($P$ < 0.05). Error bars indicate SD from three independent biological replicates, each consisting of at least 70 seedlings.
